# Predictive modeling provides insight into the clinical heterogeneity associated with *TARS1* loss-of-function mutations

**DOI:** 10.1101/2024.03.25.586600

**Authors:** Rebecca Meyer-Schuman, Allison R. Cale, Jennifer A. Pierluissi, Kira E. Jonatzke, Young N. Park, Guy M. Lenk, Stephanie N. Oprescu, Marina A. Grachtchouk, Andrzej A. Dlugosz, Asim A. Beg, Miriam H. Meisler, Anthony Antonellis

## Abstract

Aminoacyl-tRNA synthetases (ARSs) are ubiquitously expressed, essential enzymes that complete the first step of protein translation: ligation of amino acids to cognate tRNAs. Genes encoding ARSs have been implicated in myriad dominant and recessive phenotypes, the latter often affecting multiple tissues but with frequent involvement of the central and peripheral nervous system, liver, and lungs. Threonyl-tRNA synthetase (*TARS1*) encodes the enzyme that ligates threonine to tRNA^THR^ in the cytoplasm. To date, *TARS1* variants have been implicated in a recessive brittle hair phenotype. To better understand *TARS1*-related recessive phenotypes, we engineered three *TARS1* missense mutations predicted to cause a loss-of-function effect and studied these variants in yeast and worm models. This revealed two loss-of-function mutations, including one hypomorphic allele (R433H). We next used R433H to study the effects of partial loss of *TARS1* function in a compound heterozygous mouse model (R433H/null). This model presents with phenotypes reminiscent of patients with *TARS1* variants and with distinct lung and skin defects. This study expands the potential clinical heterogeneity of *TARS1*-related recessive disease, which should guide future clinical and genetic evaluations of patient populations.

**SUMMARY STATEMENT:** This study leverages an engineered, hypomorphic variant of threonyl-tRNA synthetase (*TARS1*) to capture *TARS1*-associated recessive phenotypes. This strategy revealed both known and previously unappreciated phenotypes, expanding the clinical heterogeneity associated with *TARS1* and informing future genetic and clinical evaluations of patient populations.

## INTRODUCTION

Aminoacyl-tRNA synthetases (ARS) are a family of ubiquitously expressed, essential enzymes that charge tRNA molecules with cognate amino acids, which constitutes the first step of protein translation (Antonellis and Green, 2008). The human nuclear genome encodes 37 ARS loci, with 17 encoding mitochondria-specific enzymes, 18 encoding cytoplasm-specific enzymes, and 2 encoding enzymes that function in both compartments ((Alexandrova et al., 2015; Tolkunova et al., 2000). Variants in genes encoding ARSs have been implicated in a spectrum of genetic diseases with all 37 loci implicated in recessive multisystem disorders (Meyer-Schuman and Antonellis, 2017). These disorders are caused by bi-allelic variants that severely impair gene function but do not eliminate it, as total loss of any ARS is incompatible with life. Bi-allelic pathogenic variants that affect mitochondrial ARS tend to cause phenotypes in tissues with a high metabolic demand, including leukoencephalopathies (Dallabona et al., 2014; Steenweg et al., 2012), myopathies (Sommerville et al., 2017), and liver disease (Elo et al., 2012; Walker et al., 2016). Bi-allelic pathogenic variants in ARS genes encoding cytoplasmic enzymes often affect a wider array of tissues but typically include a neurological component. The recessive neurological phenotypes associated with cytoplasmic ARSs include hypomyelination (Taft et al., 2013; Wolf et al., 2014), microcephaly (Kuo et al., 2019; Zhang et al., 2014), seizures (Salvarinova et al., 2015; Simons et al., 2015), sensorineural hearing loss, (Puffenberger et al., 2012; Santos-Cortez et al., 2013) and developmental delay (McLaughlin et al., 2010; Nowaczyk et al., 2016; Orenstein et al., 2017). Interestingly, mutations in some ARS loci cause tissue-restricted or tissue-predominant recessive phenotypes (Kuo and Antonellis, 2020). For example, although mutations in *FARSA* (Krenke et al., 2019), *FARSB* (Antonellis et al., 2018; Xu et al., 2018), *IARS1* (Orenstein et al., 2017), *MARS1* (van Meel et al., 2013), and *YARS1* (Tracewska-Siemiątkowska et al., 2017) all cause liver dysfunction as one component of a multisystem disease, only mutations in *LARS1* cause a severe, acute form of infantile liver failure (Casey et al., 2012; Casey et al., 2015; Hirata et al., 2021; Lenz et al., 2020). Similarly, pulmonary disease is pronounced in individuals with bi-allelic *FARSB* (Antonellis et al., 2018; Xu et al., 2018; Zadjali et al., 2018) and *MARS1* mutations (Rips et al., 2018; van Meel et al., 2013), including a *MARS1*-specific form of pulmonary alveolar proteinosis (Hadchouel et al., 2015). The clinical and mechanistic heterogeneities of recessive ARS-related diseases are poorly defined; advancing our knowledge in this area will require generating and characterizing relevant animal models.

Due to the conservation of ARS genes across evolutionarily diverse species, multiple model organisms can be used to study ARS biology and to investigate the impact of pathogenic variants. These models include yeast, *C. elegans*, fruit flies, and zebrafish (Malissovas et al., 2016; Oprescu et al., 2017). Mammalian models have historically been limited to studying forms of ARS-mediated dominant peripheral neuropathy (Achilli et al., 2009; Morelli et al., 2019; Seburn et al., 2006) and are less commonly employed for ARS-mediated recessive diseases, with the exception of recently reported *Dars1* (Fröhlich et al. 2021) and *Iars1* mouse models (Watanabe et al., 2023). Moving forward, mouse models will be critical tools to understand why certain tissues are particularly sensitive to loss-of-function mutations in specific ARS genes, as these questions must be addressed in a model organism with relevant tissue types.

To build a relevant model system pipeline of an understudied ARS gene, we focused on threonyl-tRNA synthetase (*TARS1*). When this study began, *TARS1* had not been implicated in any human disease phenotype. Biallelic *TARS1* variants have since been reported in two patients with a recessive brittle hair phenotype (Theil et al., 2019). To obtain a more complete assessment of *TARS1*-related recessive phenotypes in a manner that is not limited by patient ascertainment, we generated a model organism pipeline comprising yeast, worm, and mouse. We first engineered three *TARS1* missense mutations predicted to cause a loss of function and tested their effects in yeast and worm models, which revealed one of these mutations (R433H) as a hypomorphic allele. We then used R433H to study the effects of partial loss of *TARS1* function in a compound heterozygous mouse model (R433H/null). This model presents with some phenotypes that are reminiscent of the trichothiodystrophy recently attributed to bi-allelic *TARS1* mutations (Theil et al., 2019). This model also presents with distinct lung and skin phenotypes not previously associated with *TARS1*. In sum, this study expands the potential clinical heterogeneity of *TARS1*-related recessive disease, which should guide future clinical and genetic evaluations of relevant patient populations.

## RESULTS

### Identification of three loss-of-function TARS1 mutations

The model organism *Saccharomyces cerevisiae*, or Baker’s yeast, provides a tractable eukaryotic system for studying highly conserved human genes, as well as disease-associated mutations that affect the function of these genes (Kachroo et al., 2022). For aminoacyl-tRNA synthetase (ARS) genes, yeast has been a reliable model to study pathogenic mutations and test them for loss-of-function effects (Oprescu et al., 2017). Because ARS genes are highly conserved across evolution, the human open reading frame can often complement loss of the endogenous yeast ortholog, resulting in a “humanized” yeast model. In this model, yeast growth is interpreted as a proxy for ARS function, since loss of ARS function will impair cell survival and colony formation.

To design candidate recessive mutations in *TARS1*, we mutated three residues in the *TARS1* open reading frame that are highly conserved between human, mouse, worm, and yeast (Figure 1A). These mutations—N412Y, R433H, and G541R—were designed to recapitulate the types of amino-acid changes frequently observed in patient populations. To assess whether these variants affect human *TARS1* function, a complementation assay was performed using a yeast strain with the endogenous *THS1* deleted. Yeast viability was maintained with a pRS316 vector (Sikorski and Hieter, 1989) expressing the yeast *THS1*, along with *URA3.* The pYY1 vector (Chien et al., 2014) expressing either wild-type or mutant human *TARS1* was transformed into yeast, then yeast were plated on 5-FOA, which selects for the loss of the maintenance vector expressing *URA3* and *THS1* (Boeke et al., 1987). Wild-type *TARS1* supported yeast growth, demonstrating that human *TARS1* can function in yeast (Figure 1B). Transformation with N412Y *TARS1* or G541R *TARS1* did not lead to formation of colonies, indicating that these two mutations significantly impair *TARS1* function. Transformation with R433H *TARS1* did support some yeast growth but caused significantly reduced colony formation compared to wild-type *TARS1* (Figure 1B), indicating partial impairment *TARS1* function (*i.e.*, a hypomorphic allele).

**Figure 1.**
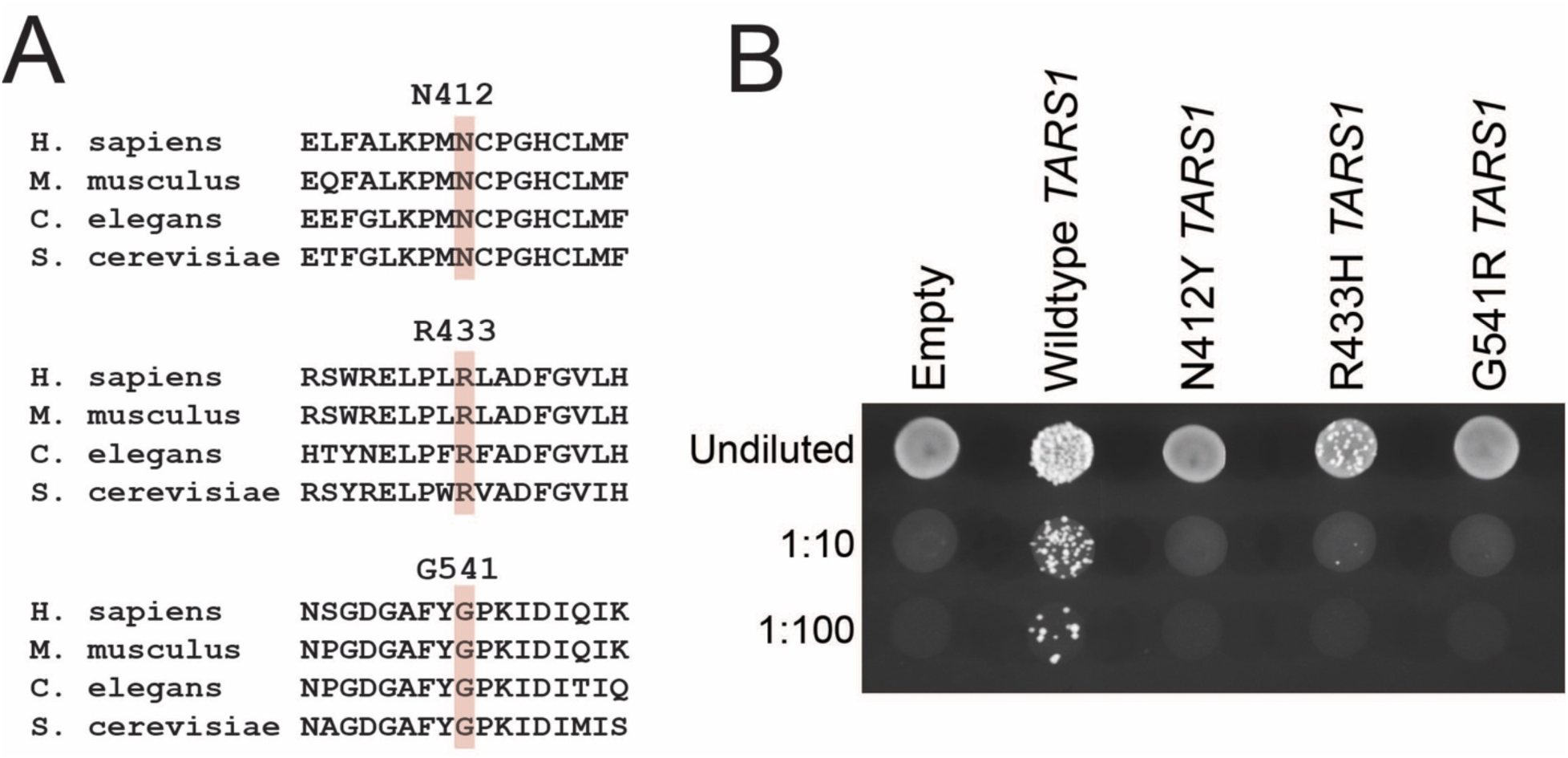
Engineered *TARS1* variants display a loss-of-function effect in yeast. **(A)** Conservation analysis of N412, R433, and G541 *TARS1*. The targeted residues are highlighted in pink, surrounded by flanking sequences from evolutionarily diverse species. **(B)** A representative image is shown from three replicates of yeast haploid strains with *THS1* deleted, and transformed with a vector with no *TARS1* insert (“Empty”), or with one to express wild-type, N412Y, R433H, or G541R *TARS1.* Yeast were spotted on media containing 5-FOA in serial dilutions (undiluted, 1:10, or 1:100) and then grown at 30°C.

### Homozygosity for G541R tars-1 is lethal in worm

To explore how loss-of-function *TARS1* variants impact the physiology of a multi-cellular organism, we modeled two of the above variants—the severe loss-of-function G541R and the partial loss-of-function R433H—in worm (*C. elegans*). The G541R or R433H mutation were each introduced into the endogenous worm *tars-1* locus using CRISPR/Cas9-mediated gene editing, with synonymous mutations introduced in *cis* to create restriction digest sites to facilitate genotyping (*EagI* for G541R and *SacI* for R433H). For simplicity, we will refer to these mutations with the human amino-acid number; however, the worm amino-acid number differs by one (Supplemental Table 1). To minimize effects from possible off-target CRISPR mutations, G541R/+ worms were back-crossed to the ancestral N2 strain five times, and R433H/+ worms were back-crossed six times. Then, to assess if either of these mutations is grossly deleterious in the homozygous state, heterozygous hermaphrodites were allowed to self-fertilize, and offspring were genotyped at the late larval L4 stage or early P1 adult stage to detect deviation from expected Mendelian ratios. In the case of the G541R/+ hermaphrodites, no G541R/G541R offspring were recovered out of 300 worms, indicating that homozygosity for G541R is lethal (Figure 2A). These data confirm that G541R is a loss-of-function allele, validating the results of the yeast complementation assay (Figure 1B). However, because the G541R/G541R genotype did not produce viable animals for additional phenotypic characterization, this mutation was not included in further studies. In contrast, when the offspring of R433H/+ hermaphrodites were genotyped, R433H/R433H homozygotes were identified. Only 33 homozygotes were identified, whereas 75 would be expected if homozygosity for R433H was benign (p<0.0001; Figure 2B). This indicates that homozygosity for R433H is deleterious but not lethal, consistent with the yeast complementation data indicating that R433H is a hypomorphic allele (Figure 1B).

**Figure 2.**
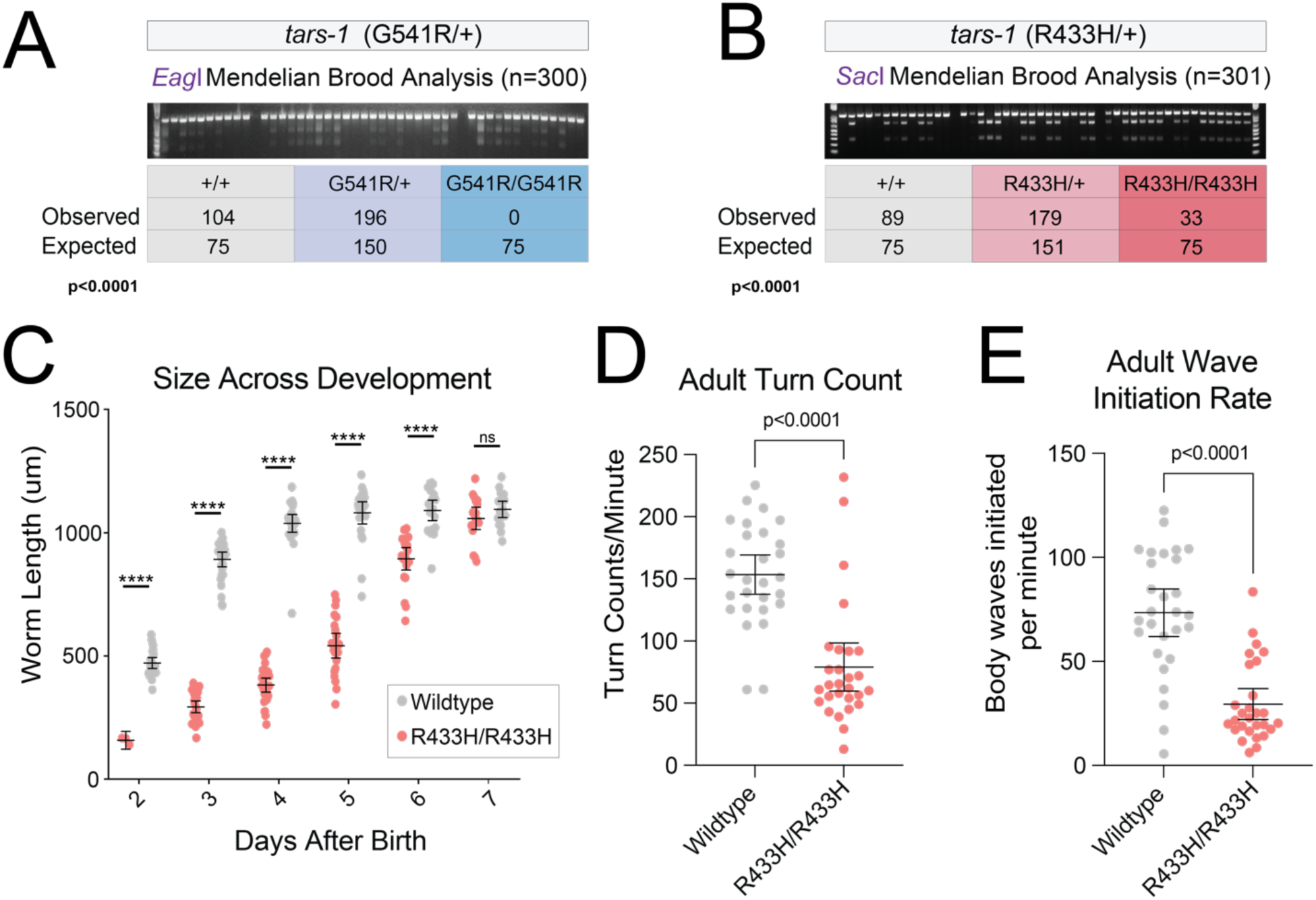
R433H *tars-1* causes reduced viability and delayed development in *C. elegans*. **(A)** Genotype analysis of offspring from five broods of G541R/+ *tars-1* hermaphrodites. A representative genotyping gel image is shown, along with the observed number of each genotype and the expected number from Mendelian segregation of a benign variant. A Chi-square test between observed and expected numbers was performed to determine statistical significance (p<0.0001). **(B)** Genotype analysis of offspring from four broods of R433H/+ hermaphrodites, exactly as described in panel A. **(C)** Measurements of body length of R433H/R433H *tars-1* worms and wild-type *tars-1* worms for six days after birth. On Day 2, n=3 for R433H/R433H, then n=18-30 worms for each subsequent day. For wild-type worms, n=18-30 for each day. **(D)** Turn count per minute for R433H/R433H worms (n=28) and wild-type worms (n=28) at adult stage P9. **(E)** Number of body waves initiated from either the head or the tail per minute, for R433H/R433H worms (n=28) and wild-type worms (n=28) at P9. For **(C-E)**, bars indicate the mean value and 95% confidence intervals. Statistical significance was evaluated using an unpaired t-test with Welch’s correction; ****, p<0.000001; ns=not significant

### R433H tars-1 causes recessive developmental delay and locomotion defects in worm

One possible explanation for the depletion of R433H/R433H *tars-1* worms in the Mendelian analysis was that this population was under-sampled compared to R433H/+ and wild-type worms. This might occur if developmental delay prevented them from reaching the genotyping timepoint at the same rate as wild-type or R433H/+ *tars-1* worms. To investigate this possibility, a population of R433H/R433H *tars-1* worms and wild-type N2 worms were age-synchronized. The physical size of the cohort was tracked for over 7 days. Beginning 48 hours after hatching, worm length was measured each day using the WormLab video and software system. R433H/R433H *tars-1* worms were consistently smaller than wild-type controls until Day 6 or 7 (Figure 2C, Supplemental Figure 1A). Whereas wild-type worms reach a mature size of approximately 1mm 3-4 days after birth, R433H/R433H *tars-1* worms do not reach this size until 6-7 days after birth.

We next investigated whether R433H might impair locomotion. We performed a thrash assay with adult worms 9 days after they reached adulthood (P9). Here, R433H/R433H *tars-*1 worms were age-matched to wild-type N2s by synchronizing embryo production. The WormLab video capture and analysis system was used to record one-minute videos of worms swimming in M9 buffer, track their motion, and calculate locomotion parameters. R433H/R433H worms display significant thrash impairment (Figure 2D and 2E; Movie 1), indicating that reduced *tars-1* function affects the neuronal circuitry or muscular function governing worm locomotion. This locomotion defect was also present in the L4 larval stage, although less pronounced (Supplemental Figure 1B and 1C). Combined with the significant delay in body size, these data indicate that R433H *tars-1* produces significant phenotypes in a multicellular eukaryotic organism.

### Partial loss of Tars1 function causes neonatal lethality in mouse due to lung defects

To determine how R433H impacts a more complex mammalian system—including defining any specific tissues that might be especially sensitive to partial loss of *TARS1* function—we developed a mouse model of R433H *TARS1.* The R433H mutation was introduced into the mouse *Tars1* locus using CRISPR-Cas9 mediated gene editing. Here, mutations are referred to with the human amino-acid number for consistency; however, the mouse amino-acid number differs by one (Supplemental Table 1). To first determine if R433H caused neonatal lethality in the homozygous state, a Mendelian analysis was performed on the offspring of a *Tars1*^R433H/+^ heterozygote intercross. Out of 43 genotyped offspring, R433H homozygotes were recovered at a frequency that did not significantly deviate from the predicted 25% (Supplemental Figure 2A), and were grossly normal throughout adulthood. These observations are consistent with our data from yeast and worm (see above) demonstrating that R433H *TARS1* retains partial function. However, it also suggests that, unlike yeast and worm, mice are less sensitive to this degree of *Tars1* impairment, and that additional reduction of *Tars1* function might be needed to observe a phenotype in a mammalian model. To this end, we generated a mouse *Tars1* null allele (F538Kfs*4) and crossed mice heterozygous for this lesion to *Tars1*^R433H/R433H^ homozygous mice. The F538Kfs*4 null allele produces a premature stop codon in exon 14, leading to a reduction in protein levels (Supplemental Figure 2C) and homozygous lethality (Supplemental Figure 2B). The R433H/F538Kfs*4 genotype resembles many individuals with recessive ARS-mediated disease who are compound heterozygous for a hypomorphic missense allele and a null allele (Meyer-Schuman and Antonellis, 2017).

To assess the effect of the R433H/F538Kfs*4 *Tars1* genotype on viability, offspring of the cross between *Tars1*^R433H/R433H^ and *Tars1*^F538Kfs*4/+^ mouse strains were genotyped at 3 weeks of age. Out of 51 pups, we only recovered 15 *Tars1*^R433H/F538Kfs*4^ (Figure 3A) offspring, indicating decreased viability prior to three weeks. An analysis of neonate deaths across four litters showed that pups that died at P0 were enriched for the R433H/F538Kfs*4 genotype; out of 15 genotyped animals presenting with neonatal death, 13 were R433H/F538Kfs*4 mice (Figure 3B). To gain insight into the neonatal pathology, a cohort of four P0 *Tars1^R433H/F538Kfs*4^* pups and three age-matched *Tars1^R433H/+^* littermates were collected for histology. The four *Tars1*^R433H/F538Kfs\**4*^ pups all died within a few hours of birth. Interestingly, one additional *Tars1^R433H/F538Kfs*4^*pup was found immediately after birth with traces of birth fluids still visible, exhibiting visibly labored breathing and a failure to right itself. This additional pup was included in the cohort to assess for a respiratory phenotype. All pups were fixed in formalin overnight and washed with 70% ethanol. Subsequently, sagittal sections were prepared and stained with H&E to detect gross morphological changes, and with Periodic Acid Schiff (PAS) staining to detect changes in glycoproteins and mucins.

**Figure 3.**
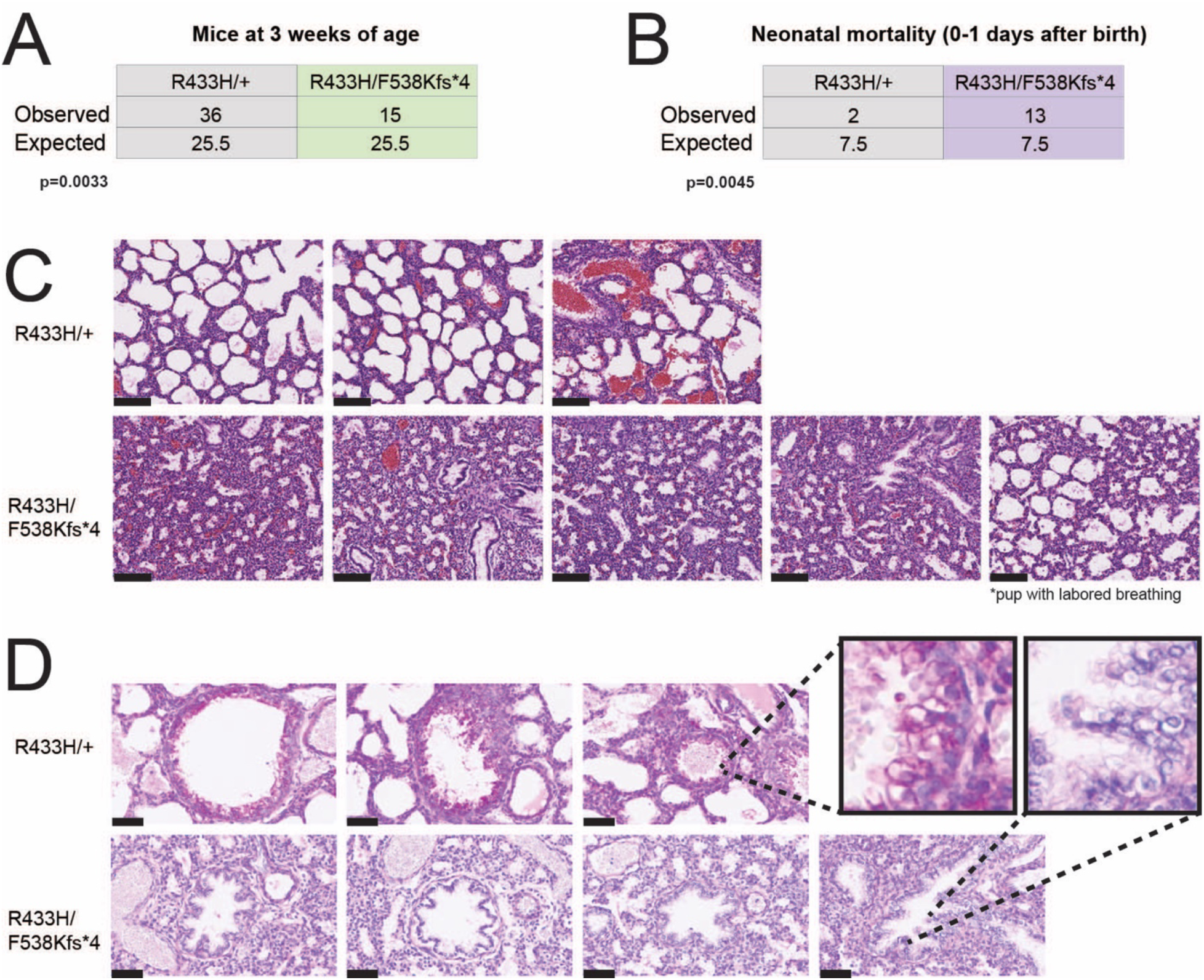
Depleted *TARS1* function causes reduced viability and a lung phenotype in a mouse model. **(A)** Genotype analysis of *Tars1*^R433H/R433H^ and *Tars1*^F538Kfs*4/+^ offspring, genotyped upon weaning at 3 weeks of age. The observed and expected number of each genotype is shown. **(B)** Genotype analysis of 15 deceased pups, identified within one day after birth. The observed and expected number of each genotype is shown. For **(A)** and **(B)**, a Chi-square test was used to determine if the difference between the number of observed and expected genotypes was statistically significant. **(C)** H&E staining of lung sections from three *Tars1*^R433H/+^ P0 pups (top row) and five *Tars1*^R433H/F538Kfs*4^ P0 pups (bottom row). All *Tars1*^R433H/+^ pups were alive when identified at P0. The first four *Tars1*^R433H/F538Kfs*4^ pups were dead at P0; the fifth was found alive with a gasping, labored breathing pattern. The black scale bar is 100um. **(D)** PAS staining of lung sections from three *Tars1*^R433H/+^ P0 pups (top row) and four *Tars1*^R433H/F538Kfs*4^ P0 pups (bottom row). Black arrows highlight the magenta PAS signal in the bronchioles of *Tars1*^R433H/+^ mice (top row), and the absence of PAS signal in the collapsed bronchioles of *Tars1*^R433H/F538Kfs*4^ mice (bottom row). The black scale bar is 50um.

The primary finding from H&E staining was an absence of air in the lungs of the four P0 *Tars1*^R433H/F538Kfs\**4*^ mice that died shortly after birth. Whereas the alveoli of *Tars1*^R433H/+^ control animals were expanded with air, the alveoli of *Tars1*^R433H/F538Kfs*^ mice were collapsed (Figure 3C). Considering the otherwise mature body development of these pups, this indicates that they died upon birth or immediately afterwards from an inability to breathe. Interestingly, the additional *Tars1*^R433H/F538Kfs*4^ pup found alive immediately after birth had only partially expanded alveoli, which correlates with the observed labored breathing. Additionally, while the bronchioles of *Tars1*^R433H/+^ control mice are replete with the magenta PAS+ signal of secretory cells, this signal is absent from the collapsed bronchioles of *Tars1*^R433H/F538Kfs*4^ animals (Figure 3D). To determine if the absent PAS+ signal indicated a loss of these secretory club cells or a significant impairment in their function, we stained similar sections with an antibody against club cell secretory protein (CCSP), an abundant lung protein primarily produced and secreted by the bronchiolar club cells in mouse (Martinu et al., 2022). This revealed no significant difference in CCSP levels between *Tars1*^R433H/F538Kfs*4^ mice and control littermates (Supplemental Figure 3), indicating that these club cells are present and grossly functional. Another possible explanation for the reduction in bronchiolar PAS+ signal was a reduction of specific threonine-rich glycoproteins like mucins, which may be poorly translated in cells with reduced *Tars1* function. Previous work in pancreatic cancer cells demonstrated that threonine starvation or knock-down of *TARS1* decreases mucin1 (MUC1) protein levels (Jeong et al., 2018). As MUC1is also a critical airway protein (Lillehoj et al., 2013), we stained our P0 pup lung sections with anti-MUC1 to determine whether decreased MUC1 levels were responsible for the decreased PAS+ signal. There was no significant difference between Muc1 signal in *Tars1*^R433H/F538Kfs*4^ pups and their littermate controls (Supplemental Figure 3). Further investigation will be required to determine the underlying mechanism of the loss of PAS+ signal and the pathophysiology of lung dysfunction in mutant *Tars1* mice.

### Partial loss of Tars1 function causes growth restriction with skin and hair abnormalities

While the studies described here were underway, a report of two patients with bi-allelic *TARS1* variants and triochothioydstrophy (TTD) was published (Theil et al., 2019). The phenotypes described in these two patients included delayed physical development, ichthyosis, and collodion baby, and the brittle hair of TTD. Interestingly, *Tars1*^R433H/F538Kfs*4^ mice display phenotypes that are reminiscent of these human disease features. For example, *Tars1*^R433H/F538Kfs*4^ mice that survived to adulthood were, on average, smaller than their *Tars1*^R433H/+^ littermates (Figure 4). Reduced body weight was more consistent in females (Figure 4B) than males (Figure 4C), who reach a normal body size by seven weeks of age. This reduced size is consistent with the delayed physical development described in *TARS1* patients (Theil et al., 2019), and with the growth restriction phenotypes in patients with other ARS-mediated recessive disease (Kuo et al., 2019; Orenstein et al., 2017; Peroutka et al., 2019; Zadjali et al., 2018).

**Figure 4.**
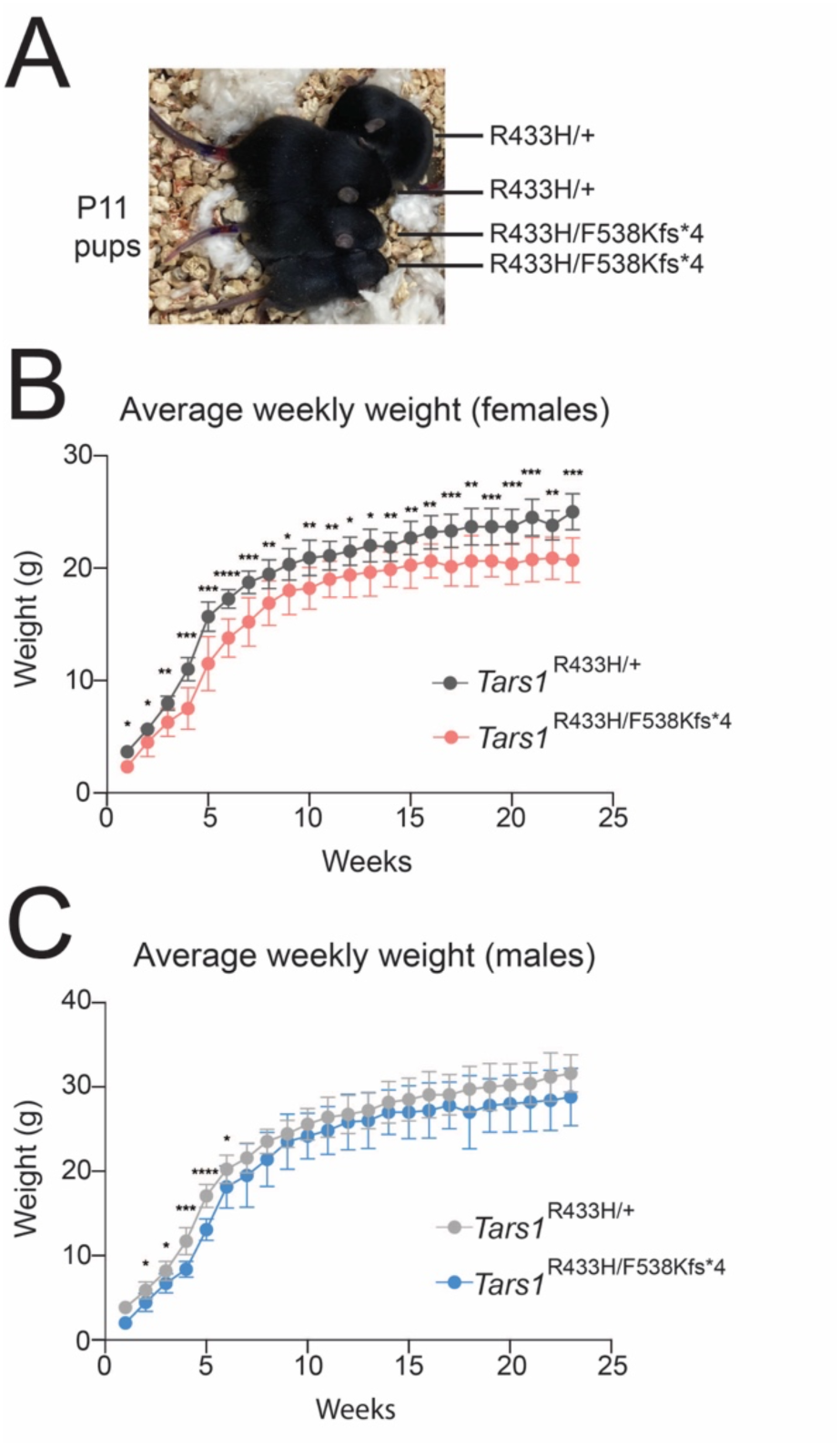
Depleted *Tars1* function causes reduced body size in a mouse model. **(A)** Image of four littermates at P11, grouped together for comparison of body size. The genotype of each mouse is provided. **(B)** The average weekly weights of female *Tars1*^R433H/F538Kfs*4^ mice (n=9) and female *Tars1*^R433H/+^ (n=11) littermates are shown, until 23 weeks of age. **(C)** The average weekly weights of male *Tars1*^R433H/F538Kfs*4^ mice (n=6) and male *Tars1*^R433H/+^ (n=12) littermates are shown, until 23 weeks of age. For **(B)** and **(C)**, bars represent the mean value and one standard deviation. An unpaired t-test was performed for each week to determine if the difference between the two genotypes was statistically significant. **** p<0.0001, *** p<0.001, ** p<0.01, * p<0.05. All values in **(C)** that are not marked with an asterisk are not significantly different.

We also detected skin and hair abnormalities in the *Tars^R433H/F538Kfs*4^*P0 pups and adult mice. The pups had a thinner epidermal layer than control littermates, with fewer layers of stratum corneum (Figure 5A and 5B). They also displayed variable degrees of hair follicle hypoplasia (Figure 5A). This is unlike the classic mouse model of TTD, which exhibits a thicker epidermal layer (de Boer et al., 1998). Adult *Tars1*^R433H/F538Kfs*4^ mice also displayed a striking postnatal hair phenotype, although it did not resemble the sparse, brittle hair associated with TTD, and was not as severe as the hair loss previously described in the TTD mouse model (de Boer et al., 1998). In a nine-litter cohort, 10 out of 14 *Tars1*^R433H/F538Kfs*4^ mice (71.4%) lost hair on their heads and/or upper back by 23 weeks of age, compared to 1 out of 23 *Tars1*^R433H/+^ littermates (4.4%). Hair loss onset occurred between 13 and 23 weeks of age (Figure 6A) and followed a stereotypic pattern of bald spots on the head and/or along the scapula of the upper back (Figure 6B). In more advanced stages, it spanned the entire upper back (Figure 6C), although it did not encompass the majority of the body as previously described for TTD mice, nor did it grow back in cycles of loss and regrowth (de Boer et al., 1998). To more thoroughly define this phenotype, histopathology was performed on hair samples from the affected regions for three *Tars1*^R433H/F538Kfs*4^ mice and three *Tars1*^R433H/+^ littermates; one pair was 2 months old, another was 12 months old, and the third was 14 months old. Analysis of H&E staining did not reveal gross abnormalities in hair follicles (data not shown), although this analysis was complicated by the asynchronous hair cycling of adult mice. We also did not observe the classic “tiger tail banding pattern” seen under polarizing microscopy hair from TTD patients (Faghri et al., 2008). Taken together, our data demonstrate that partial loss of *TARS1* function causes unusual hair and skin phenotypes in mouse, impairs body weight, and causes a partially penetrant but severe respiratory deficiency.

**Figure 5.**
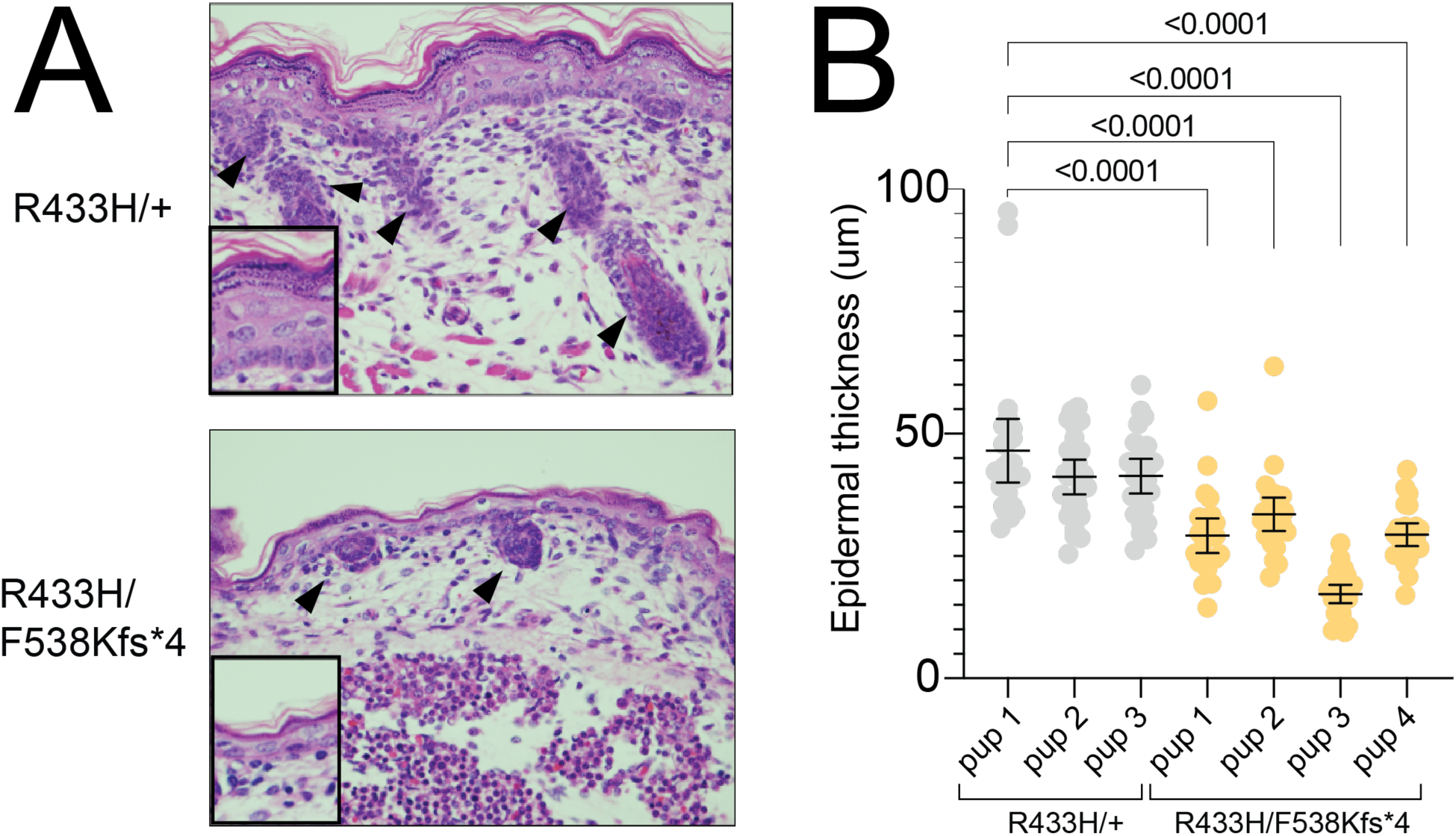
Depleted *Tars1* function causes skin phenotypes in a mouse model. **(A)** H&E staining of dorsal skin sections from P0 pups. The upper panel shows a representative image of skin from a *Tars1*^R433H/+^ mouse, and the bottom panel shows a representative image of skin from a *Tars1*^R433H/F538Kfs*4^ mouse. Black arrows point to hair follicles in each image. **(B)** Measurements of epidermal thickness on four *Tars1*^R433H/F538Kfs*4^ P0 Pups and three *Tars1*^R433H/+^ P0 littermates (n=25 measurements per pup). Bars indicate the mean value and 95% confidence interval. Statistical significance was determined with a one-way ANOVA with Šidák’s multiple comparisons testing, comparing all animals to R433H/+ pup 1. Only p values p<0.05 are shown (differences between R433H/+ pups were not statistically significant).

**Figure 6.**
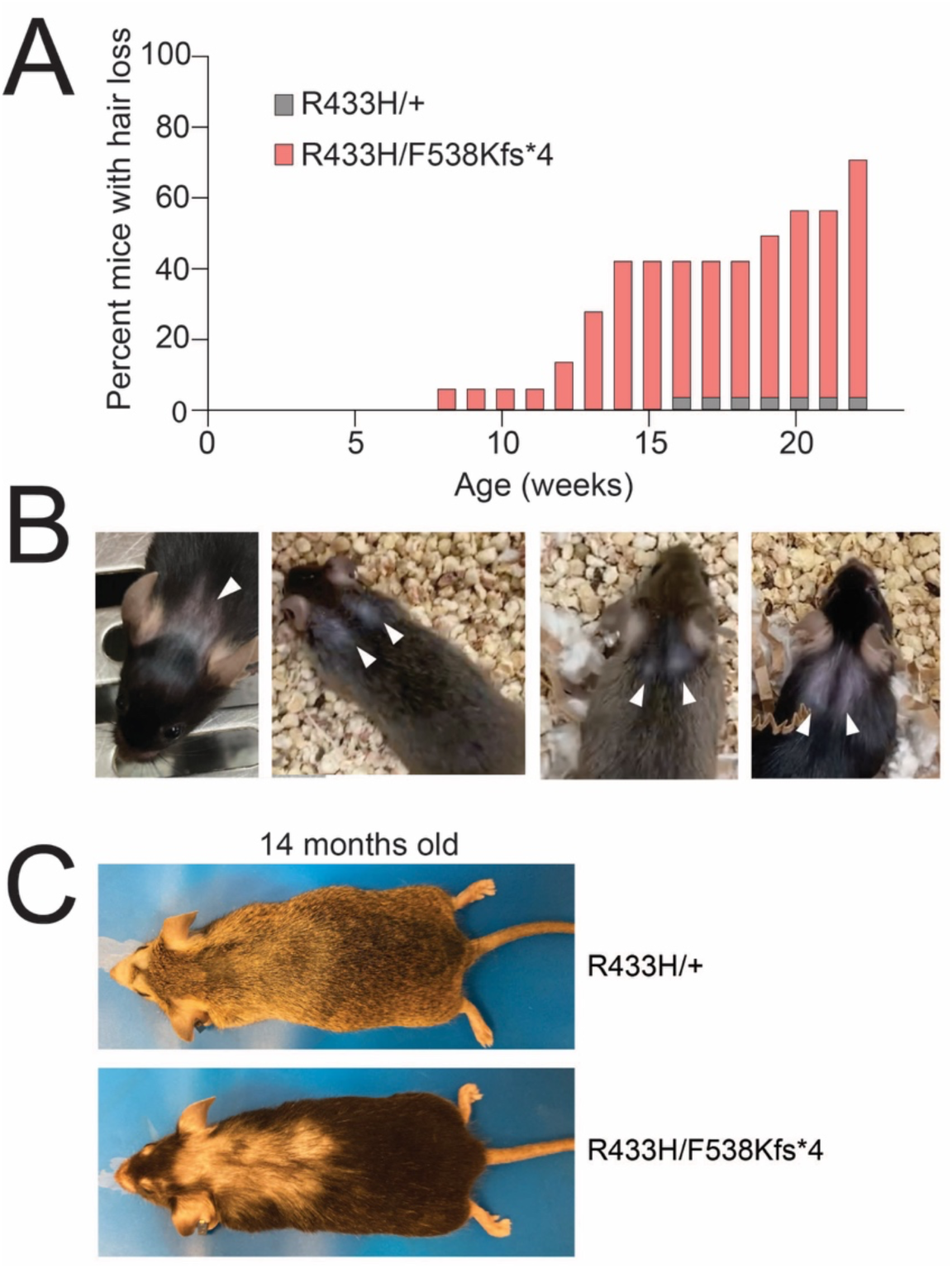
Depletion of *Tars1* function causes hair loss in a mouse model. **(A)** The cumulative percentage of *Tars1*^R433H/F538Kfs*4^ mice (pink, n=14) and *Tars1*^R433H/+^ mice (gray, n=23) with hair loss on the back of the head or upper back is shown, until 23 weeks of age. **(B)** Representative images of four individual *Tars1*^R433H/F538Kfs*4^ mice with hair loss; white arrows point to the consistent pattern of upper back bald patches. The depicted mice are between 10 weeks and 17 weeks of age. **(C)** A representative image of hair phenotypes in *Tars1*^R433H/+^ (top) and *Tars1*^R433H/F538Kfs*4^ (bottom) animals. Note that extended hair loss stretches from the head to the middle of the back in the *Tars1*^R433H/F538Kfs*4^ mouse, at 14 months of age. A *Tars1*^R433H/+^ littermate is shown above, with signs of barbering by the nose and mild age-related hair thinning on the back.

## DISCUSSION

In this study, we leveraged the established characteristics of pathogenic ARS variants to develop a model system pipeline for predicting the clinical heterogeneity of *TARS1*-related recessive disease. We successfully engineered two loss-of-function *TARS1* missense mutations, including a partial loss-of-function allele, R433H, that we employed to explore recessive phenotypes across three model organisms. This analysis revealed that R433H: (**1**) partially reduced yeast growth in yeast complementation assays; (**2**) caused developmental delay and locomotion defects in homozygous worms; and (**3**) caused lung failure, decreased body size, and skin and hair defects in mice when modeled *in trans* with a null allele in mice. While skin and hair phenotypes are seen in our mouse mutants, they do not precisely mimic those recently described for humans with *TARS1* mutations. The remaining phenotypes are unique and have not been previously described. Importantly, our data indicate that phenotypic heterogeneity will ultimately be observed in human *TARS1*-related recessive disease, and that individuals with bi-allelic *TARS1* variants should be carefully evaluated for lung disease.

It is interesting to consider why some tissues may be particularly sensitive to reductions in *TARS1* function. One possibility is that critical proteins with a particularly high threonine content, such as mucins, are more dramatically affected by decreased *Tars1* activity. This could lead to defects in the tissues that rely heavily on these proteins, such as the lung. Interestingly, the gut is also dependent on mucin synthesis (Wagner et al., 2018). Although preliminary investigation of gut histology in P0 mice did not identify any changes in PAS signal, careful assessments for gut phenotypes should be performed on patients that are homozygous or compound heterozygous for pathogenic *TARS1* variants. Another possibility is that decreased *Tars1* activity reduces the available population of charged tRNA^Thr^, triggering eIF2α phosphorylation which then leads to decreased global protein translation. This might affect cells with a high demand for protein translation, such as transient-amplifying progeny of stem cells. For example, if aging hair follicle stem cell progeny cannot properly translate the large mass of proteins required for differentiation of the multiple cell types comprising the mature hair follicle and hair shaft, this could explain a failure to regrow hair in the adult *Tars1*^R433H/F538Kfs*4^.

In summary, this study demonstrates the efficacy of using variant engineering and a tiered model organism approach to predict the pathogenicity of variants in ARS genes. While additional research on human subjects and animal models will be required to fully define the clinical heterogeneity of *TARS1*-related disease, the data presented here should be useful in genetic and clinical evaluation of relevant patient populations.

## MATERIALS AND METHODS

### Generation of TARS1 expression constructs

The open reading frame (ORF) of *TARS1* was amplified from HeLa cell cDNA, using primers with the attB1 and attB2 gateway recombination sequences (primer sequences in Supplemental Table 2). These amplicons were purified with Qiagen Spin Miniprep columns and recombined into pDONR221 using Gateway cloning technology (Invitrogen). The recombination reaction was then transformed into Top10 cells (Invitrogen) to isolate clonal populations. Individual bacterial colonies were selected and grown in media containing kanamycin, which selected for the kanamycin resistance cassette on pDONR221. Plasmids were then isolated using the Qiagen Miniprep kit and genotyped by digesting with *Bsr*GI (New England Biolabs) to detect the presence of the *TARS1* insert. Clones with successful insertions were analyzed by Sanger sequencing to ensure absence of mutations introduced by amplification errors. To introduce variants into the *TARS1* ORF, site-directed mutagenesis was performed with the QuickChange II XL Site-Directed Mutagenesis Kit (Agilent) (primer sequences Supplemental Table 2). The reaction was transformed into Top10 cells and grown in LB containing kanamycin to select for pDONR221. Plasmid DNA was isolated and sequenced as above, to ensure successful mutagenesis. Then, the Gateway LR reaction was used to recombine the wild-type or mutant *TARS1* into the vector pYY1. This vector has a 2-micron origin of replication, resulting in a high copy number per cell, as well as the *ADH1* promoter, resulting in strong constitutive *TARS1* expression. Recombinants were transformed into Top10 cells, which were plated on ampicillin to select for the ampicillin resistance cassette on pYY1. Plasmids were extracted, purified, and digested with *Bsr*GI to identify successfully recombined clones.

### Yeast complementation assays

Yeast complementation assays were performed with the Δ*THS1* strain (Horizon Discovery, Clone ID 21471). Yeast viability was maintained with a pRS316 vector that expresses wild-type *THS1* from the endogenous yeast promoter. pRS316 also carries the auxotrophic marker *URA3*, and has a yeast centromere sequence which results in a low copy number per cell. The pYY1 vector (expressing wild-type *TARS1*, mutant *TARS1*, or an empty control) was transformed into yeast with a standard lithium acetate transformation, performed at 30°C with 200ng of plasmid. Yeast were grown on solid media without uracil and leucine, which selected for cells with both pRS316 and pYY1. Yeast were grown for 3 days at 30°C, then individual colonies were picked into 2mL liquid media lacking uracil and leucine. These cultures were grown for 2 days at 30°C, shaking at 275 rpm. Then, 1mL of saturated culture was centrifuged at 15,000 rpm for 1 minute and cell pellets were re-suspended in 50μl water. Yeast were serially diluted to 1:10, 1:100, or 1:1000 using water. 10μl of each dilution (included undiluted yeast) was spotted on complete media containing 5-FOA (Teknova), which selects for cells that have spontaneously lost the pRS315 vector expressing *URA3* and *THS1* (Boeke et al., 1987). After 3 to 5 days, yeast growth was visually inspected.

### Generation of the G541R and R433H tars-1 C. elegans strains

To generate the G541R and R433H *tars-1* models, CRISPR-Cas9 genome editing was performed according to previously described methods (Prior et al., 2017). Briefly, the gonadal tract of P1 adult worms was injected with an injection mix of: 300mM KCl, 20mM HEPES, 2.5 ng/μl pCFJ90, 50ng/μl single stranded oligonucleotide homologous donor repair template (Integrated DNA Technologies), 5μM single guide (sg) RNA (Synthego), and 5μM Cas9 protein (Integrated DNA Technologies). Sequences for the repair templates and guide RNAs can be found in Supplemental Table 3. Injected worms were then placed on single 35mm plates of nematode growth media (NGM) and fresh OP50 bacteria as a food source. Approximately 2 days after injection, plates were screened for the presence of F1 progeny expressing the pCFJ90 marker, which expresses mCherry in the pharyngeal muscles. This enriches for worms that were exposed to the injection mix, increasing the likelihood of identifying a worm subjected to genome editing. The mCherry-positive F1s were singled to individual plates and allowed to produce their own offspring (F2). Then, the F1 worms were placed in lysis buffer (50mM KCl, 10mM Tris-HCl pH 8.3, 2.5mM MgCl2, 0.45% NP-40, 0.45% Tween-20, 1mg/mL proteinase K) and lysed with incubation at -80°C for one hour, incubation at 65°C for one hour, and incubation at 95°C for fifteen minutes. To genotype worms, the targeted *tars-1* region was amplified by PCR (primer sequences in Supplemental Table 2) using Q5 PCR mix (New England Biolabs). Amplicons were then purified with DNA Clean and Concentrator kits (Zymo Research) and digested with the appropriate restriction enzyme (*Eag*I for G541R or *Sac*I for R433H, New England Biolabs). Digested PCR products were separated on a 1% agarose gel and analyzed to identify successful integration of the restriction site. The undigested PCR product from F1s with successful gene editing events was submitted for Sanger sequencing to confirm proper insertion of the restriction site and the desired *tars-1* mutation. The offspring of these F1 worms were then maintained for subsequent experiments. To reduce possible off-target mutations caused by CRISPR-Cas9 editing, R433H/+ *tars-1* worms were back-crossed to the ancestral N2 strain six times. To assess the Mendelian ratios of the offspring of R433H/+ *tars-1* worms, 6-8 worms from the R433H/+ strain were singled to individual 35mm plates with OP50, allowed to self-fertilize and produce progeny, then genotyped to confirm heterozygosity for R433H. After confirmation, individual progeny were picked into wells of a 96 well plate for genotyping and Mendelian ratio analysis. This was repeated four times for a total of 301 genotyped offspring.

### Measuring worm body size through development

To identify differences in rates of development, R433H/R433H *tars-1* worms and wild-type N2 worms were first age-synchronized by placing approximately 25 adult worms on a 60 mm plate with NGM and OP50, letting them produce embryos for 4-5 hours, and then removing the adults. After 48 hours, worms were transferred to unseeded 35mm NGM plates in batches of 4-5 worms. These worms were filmed and analyzed using the WormLab System (MBF Biosciences). Plates were filmed for 30-second intervals, with the camera set at 4.81um/pixel for R433H/R433H worms (Setting 1 on the Wormlab camera apparatus [MBF Bioscience]) and 8.47um/pixel for N2 worms (Setting 3 on the Wormlab camera apparatus [MBF Bioscience]). After filming, worms were moved to new NGM plates seeded with OP50. Filming was repeated every 24 hours up to 168 hours, or 7 days, after birth (as R433H/R433H worms increased in size, filming was performed with the camera setting at 8.47um/pixel). All videos were analyzed with the WormLab software (MBF Bioscience), and the ‘worm length’ parameter was extracted to compare the size of R433H/R433H *tars-1* worms and N2 worms over the course of development.

### Worm thrash assays

Thrash assays were performed to detect changes in worm movement. The bottom of each well of a Nunc 4-well dish (Thermo Scientific) was coated with 2.5% agarose. 500μl liquid M9 media (22mM H2KO4P, 42 mM HNa2O4P, 85 mM NaCl, 1mM MgSO4) was added to each well, and 1-5 worms (wild-type or R433H/R433H *tars-1*) were placed in the M9. Worms were allowed to acclimate for 30-60 seconds before they were filmed with the WormLab System (MBF Biosciences) for 1 minute. Only worms with at least 1,000 frames or 30 seconds of high-quality video were included in subsequent analysis. To identify defects in locomotion, the WormLab parameters “Turn count” and “Wave initiation rate” were analyzed.

### Generation of Tars1 mouse lines

The R433H mutation was introduced into the mouse *Tars1* locus using CRISPR-Cas9 mediated gene editing, which was performed by the University of Michigan Transgenic Animal Core. A single-stranded oligonucleotide (ssODN) was designed to introduce the R433H mutation *in cis* with silent mutations that ablated a *Bgl*I cut site that is present in the wild-type allele, and prevented binding of the guide RNA after repair. Cas9, sgRNA, and ssODN were injected into hybrid C57BL/6J x SJL/J F1 zygotes, which were implanted into pseudopregnant females. These mice produced 32 pups, which were genotyped by PCR-amplification (primer sequences in Supplemental Table 2) and *Bgl*I digestion to identify mice that had incorporated the repair template. Amplicons were submitted for Sanger sequencing to identify mice with proper integration of the repair template. These mice were mated to C57BL/6 mice to establish germline transmission. To assess the Mendelian ratios of offspring genotypes, 43 pups from crosses between *Tars1*^R433H/+^ females and *Tars1*^R433H/+^ males were genotyped using the *Bgl*I restriction enzyme digest strategy described above.

The F538Kfs*4 allele was similarly generated, using a sgRNA that targeted exon 14. This generated an 11 base pair deletion that ablates a *Hae*III cut site and lead to a premature stop codon shortly downstream of the PAM site (F538Kfs*4). To genotype this deletion, the region was amplified (primer sequences in Supplemental Table 2) and digested with *Hae*III; an undigested upper band indicated the presence of the frameshift allele. To evaluate the Mendelian ratios of offspring genotypes, 28 pups from crosses between *Tars1*^F538Kfs*4/+^ females and *Tars1*^F538Kfs*4/+^ males were genotyped using this restriction enzyme digest strategy.

### Western blot analyses from mouse brain

Total protein concentration was measured using the Thermo Scientific Pierce BCA Protein Assay Kit. 6.25μg, 12.5μg, or 25μg of lysate was analyzed. Samples were prepared with 1X Novex Tris-Glycine SDS sample buffer (Invitrogen) and 2-mercaptoethanol (BME), and were boiled at 99°C for 5 minutes. Protein samples were separated on precast 4-20% Novex Wedgewell Tris-glycine gels (Invitrogen) at 150V for 1 hour and 15 minutes. PVDF membranes (Millipore Sigma) were pre-washed in 100% methanol for 1 minute, then soaked in 1X transfer buffer (Invitrogen) and 10% methanol between two pieces of filter paper (Thermo Fisher Scientific). Samples were transferred from the Tris-Glycine gel to the PVDF membrane using a Mini Trans-Blot Electrophoretic Transfer Cell (Biorad) at 0.03A for 18-20 hours. The membrane was blocked in 2% milk in 1X TBST overnight at 4°C. After blocking, the membrane was washed with 1X TBST three times, with each wash comprising five minutes of rocking at room temperature. Primary antibody was applied in a 2% milk solution: anti-TARS1 (Thermo Fisher PA5-30690) was applied at 1:500 dilution and anti-actin (Sigma A5060) was applied as a loading control at 1:5,000 dilution. Primary antibody was incubated overnight at 4°C. Membranes were washed three times with 1X TBST as above, and secondary antibodies (anti-mouse HRP [1:2,000; Thermo Fisher Scientific] for TARS1 and anti-rabbit HRP [1:5,000; EMD Millipore] for actin) were applied in 2% milk solution. The blots were rocked for 1 hour at room temperature before incubating with SuperSignal West Dura substrate (Thermo Scientific) according to the manufacturer’s instructions.

### Preparation of mouse tissues for histology

To investigate P0 pups for gross histological changes, dead pups were collected and live pups were sacrificed by decapitation. Pups were individually fixed in neutral-buffered formalin, rocking overnight at room temperature. Pups were then placed in 70% ethanol and stored at 4°C. To investigate histological changes in the hair and skin of adult mice, three *Tars1*^R433H/F538Kfs*4^ mice and their age-matched, sex-matched *Tars1*^R433H/+^ littermates were sacrificed. The mice were shaved, and skin was collected from the dorsal trunk, ventral trunk, ears, tail, and paws, as well as from the area of the head with visible hair loss. Skin samples were placed on 0.45μm HA filters (Millipore) wetted in PBS and strips were cut parallel to the direction of hair follicle growth. All strips were then fixed overnight in neutral-buffered formalin at room temperature, transferred to 70% ethanol and stored at 4°C. Samples were shipped to Histoserv, Inc. for embedding and sectioning. Briefly, samples with bone were decalcified, tissues were dehydrated, and water inside of the tissues was replaced with paraffin wax. Tissues were then embedded into wax blocks of paraffin. Blocks were sectioned and affixed to slides (two sagittal sections were taken for the P0 pups). Adult skin sections were stained with H&E; P0 pup sections were stained with either H&E or PAS, which detects glycoproteins and mucins.

### Immunohistochemistry

Paraffin sections were deparaffinized and rehydrated through a graded ethanol series. Endogenous peroxidase activity was quenched by treatment with Peroxidase Suppressor Reagent for 15 minutes (Pierce™ Peroxidase IHC Detection Kit, Thermos Scientific, 36000). Heat induced epitope retrieval was performed by incubating slides in antigen retrieval agent (R&D Systems, CTS015) at 95°C for 5 minutes and then allowing the slides to cool at room temperature for 10 minutes. Slides were blocked in Universal Blocker Blocking Buffer in TBS (Pierce™ Peroxidase IHC Detection Kit, Thermo Scientific, 36000) for one hour. Primary antibodies (see below) were diluted in blocking buffer and slides were incubated with primary antibody overnight at 4°C. After washing with PBS, slides were incubated in secondary antibody for one hour at room temperature. Substrate incubation, chromogenic development, and hematoxylin counterstain were performed according to manufacturer recommendation (Pierce™ Peroxidase IHC Detection Kit, Thermo Scientific, 36000). Primary antibodies used were rabbit anti-CCSP (Sigma-Aldrich, 07-623) 1:1000 and rabbit anti-MUC1 (Abcam, ab15481) 1:100. For anti-MUC1 staining, secondary antibody (Sigma Aldrich, 12-348) was used at a dilution of 1:500; for anti-CCSP staining, secondary antibody was used at a dilution of 1:1000.

### Analysis of epidermal thickness in P0 pups

Dorsal skin from H&E-stained sections was used to analyze the epidermal thickness of four *Tars1*^R433H/F538Kfs*4^ mice and three *Tars1*^R433H/+^ littermates. Five 1mm areas were selected, evenly spaced out across the back. In each 1 mm area, the thickness of the epidermis was measured by drawing lines in Adobe Illustrator that span the width of the epidermal layer, then using the 200um scale in each image to convert line length to um. Five measurements were made that evenly spanned the 1mm area; each measurement was made at the widest local area.

## ACKNOWLEDGEMENTS

The authors would like to thank Dr. Thomas Saunders and University of Michigan Transgenic Animal Model Core, and Dr. Sunny Wong for his advice on the skin phenotypes of the *Tars* mouse model.

## DATA AVAILABILITY

No large datasets were generated in the course of this study. All raw data files are available upon request.

## AUTHOR CONTRIBUTIONS

Conceptualization: R.M-S., A.A, A.B.. Methodology: R.M-S., S.O., Y.P., G.L., M.G., A.D., A.B., M.M., A.A. Validation: A.C, K.J.. Investigation: R.M-S., J.P., S.O., M.G. Resources: A.A, A.B., M.M. Writing: R.M-S., A.A. Funding acquisition: A.A., M.M.

## SUPPLEMENTAL MATERIALS

**Movie 1. R433H/R433H worms display significant locomotion defects.** A ten-second video of adult (P9) wildtype worms (N2 strain) thrashing in liquid M9 media, followed by a ten-second video clip of P9 R433H/R433H *tars-1* worms thrashing in liquid M9.

(not included in preprint)

---

**Supplemental Figure 1.**
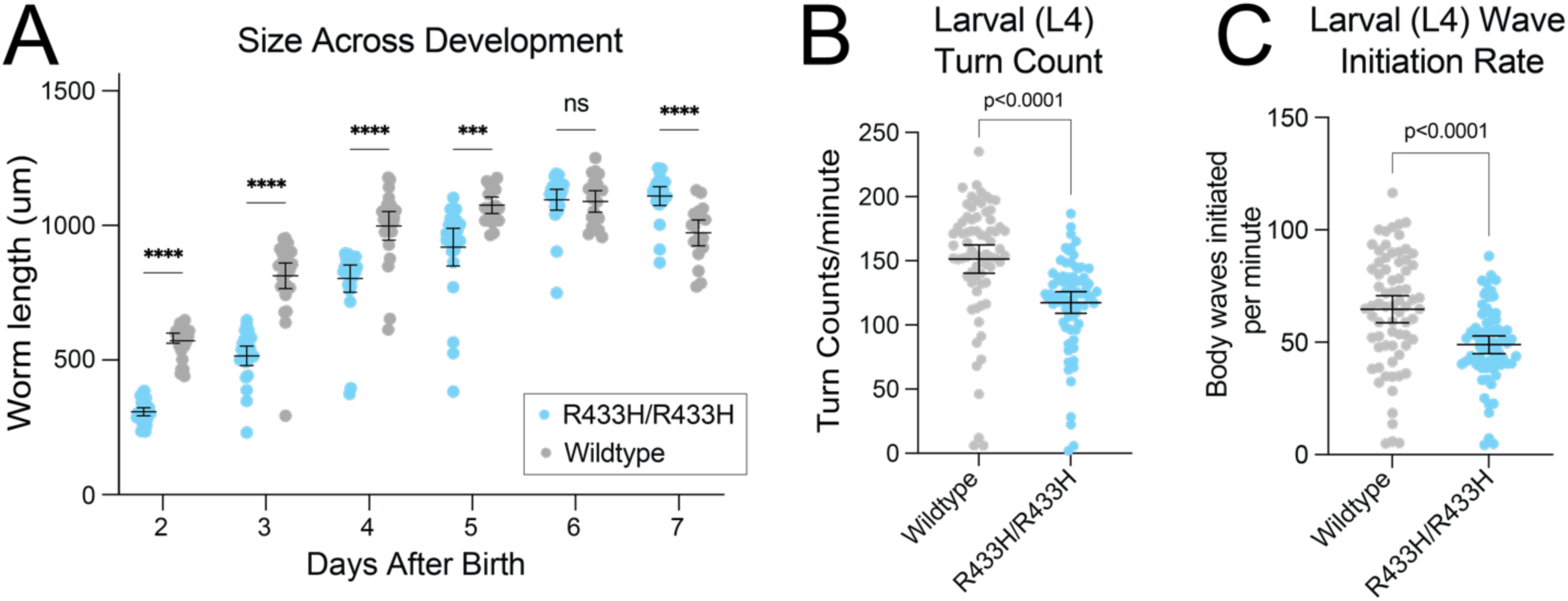
**(A)** Body length measurements of R433H/R433H *tars-1* worms and wild-type *tars-1* worms from two to six days after birth. For R433H/R433H, 24-29 worms were measured each day. For wild-type worms, 19-35 were measured each day. **(B)** Turn counts per minute for R433H/R433H worms (n=73) and wild-type worms (n=71) at larval stage L4, which was identified based on gonadal development. **(C)** Rate of wave initiations from either the head or the tail for L4 R433H/R433H worms (n=73) and wild-type worms (n=71). For all panels, bars indicate mean value and 95% confidence intervals. Statistical significance was evaluated using an unpaired t-test with Welch’s correction; ****, p<0.0001; ***, p<0.001 ns=not significant.

**Supplemental Figure 2.**
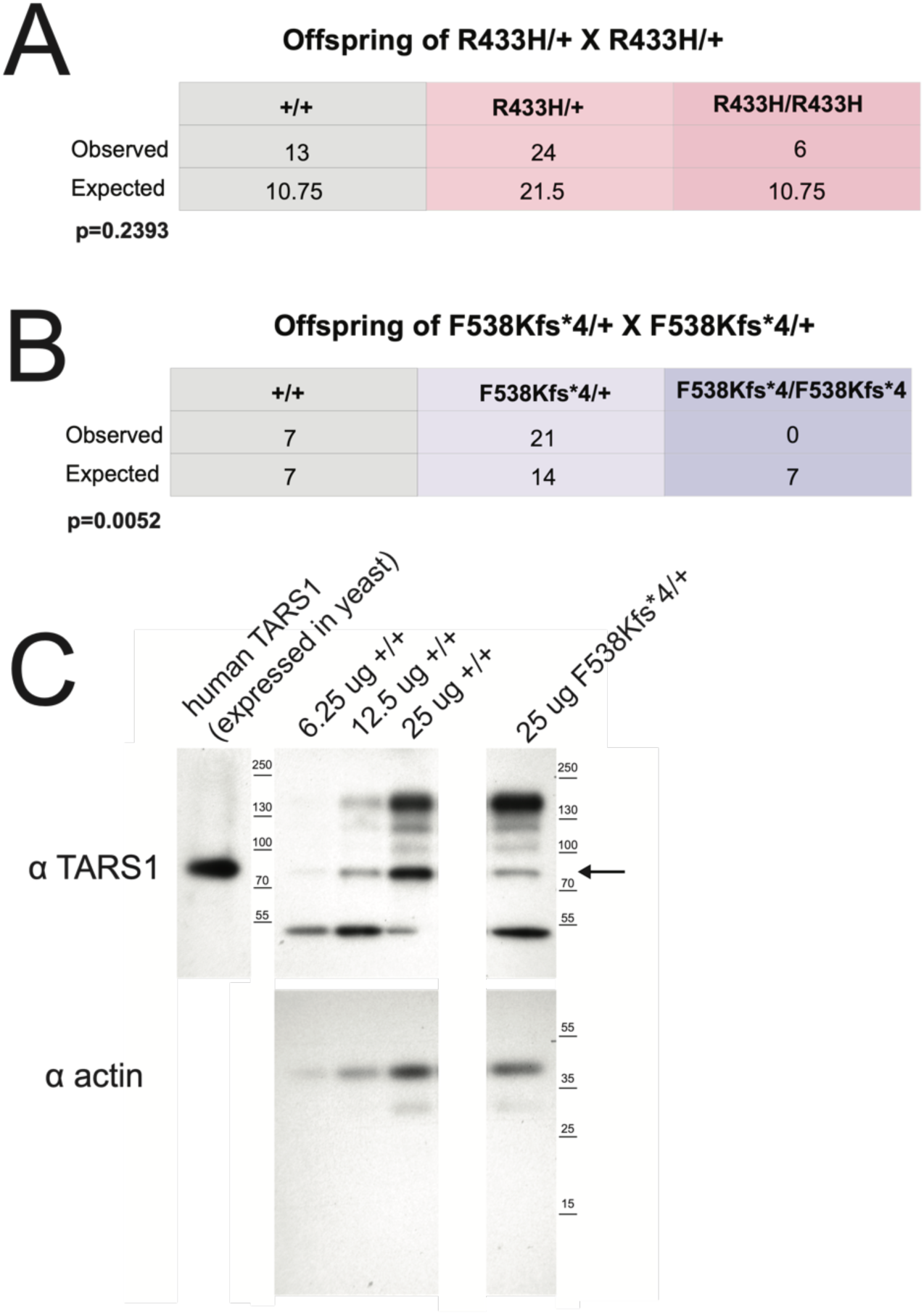
**(A)** Genotype analysis of 28 offspring from *Tars1*^F538Kfs*4/+^ x *Tars1*^F538Kfs*4/+^ mouse mating pairs. **(B)** Genotype analysis of 43 offspring from *Tars1*^R433H/+^ x *Tars1*^R433H/+^ mouse mating pairs. All mice were genotyped at approximately 3 weeks of age. Chi-square tests were performed to determine if the difference between observed genotype counts and expected genotype counts was statistically significant. **(C)** Representative western blot image for Tars1 protein in brain tissue of wild-type *Tars1* mice and *Tars1*^F538Kfs*4/+^ mice. Human TARS1 (predicted size of 83 kDa), expressed in yeast, is shown on the left as a size control. An arrow points to the band corresponding to Tars1 (predicted size of 83 kDa). For wild-type mice, 6.25μg, 12.5μg, and 25μg lysate was loaded as a comparison to 25μg lysate for *Tars1*^F538Kfs*4/+^ samples.

**Supplemental Figure 3.**
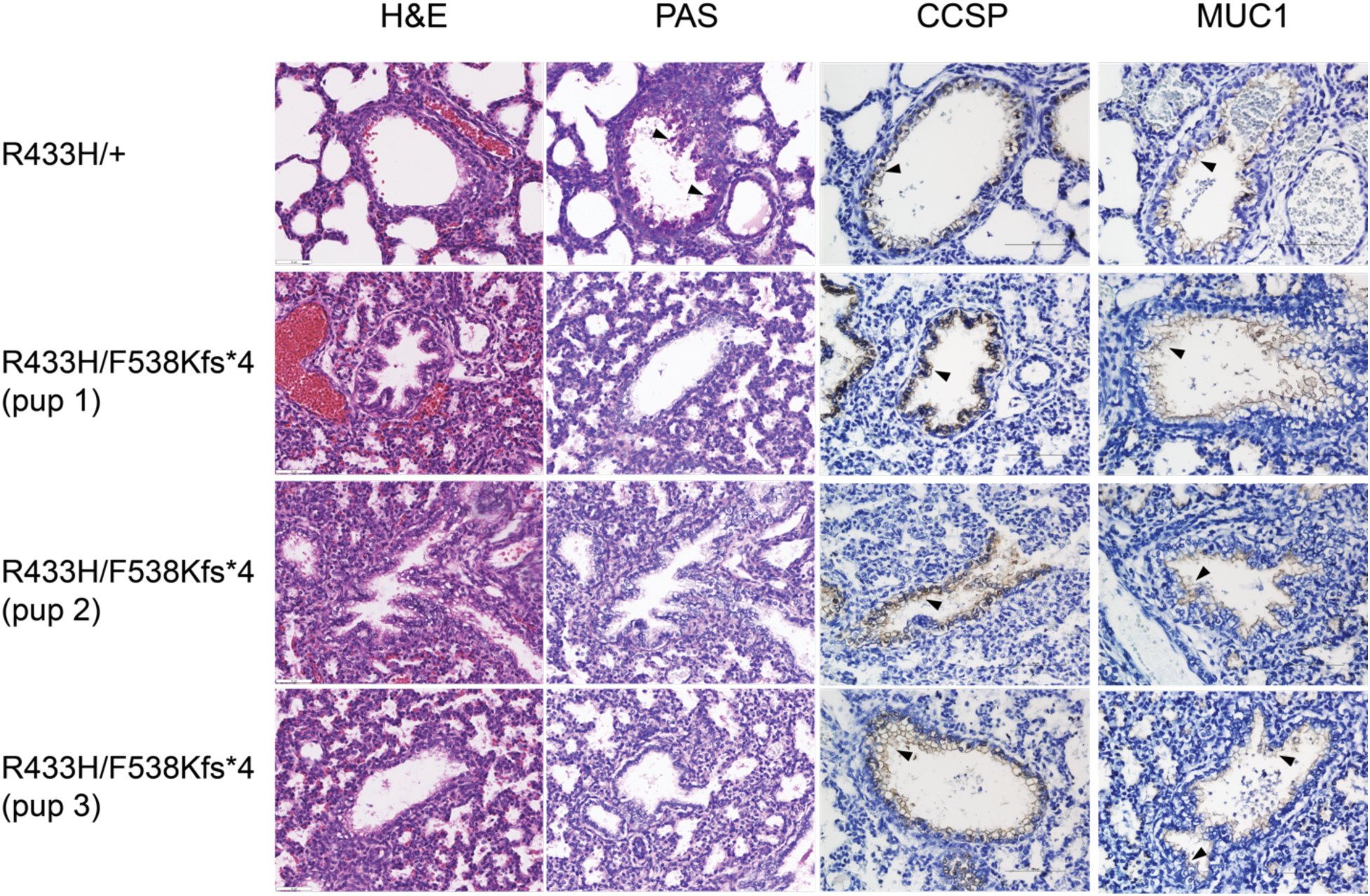
Lung sections stained with (from left to right) H&E, PAS, CCSP antibody, or MUC1 antibody. The R433H/+ images (top row) are representative of three R433H/+ P0 pups, and the R433H/F538Kfs*4 images (bottom three rows) are representative of five R433H/F538Kfs*4 P0 pups. Black arrows point to the PAS signal in the R433H/+ mouse, and the CCSP and MUC1 signals in all mice. The CCSP and MUC1 signals are shown in brown, with the hematoxylin counterstain in dark blue.

**Supplemental Table 1.**
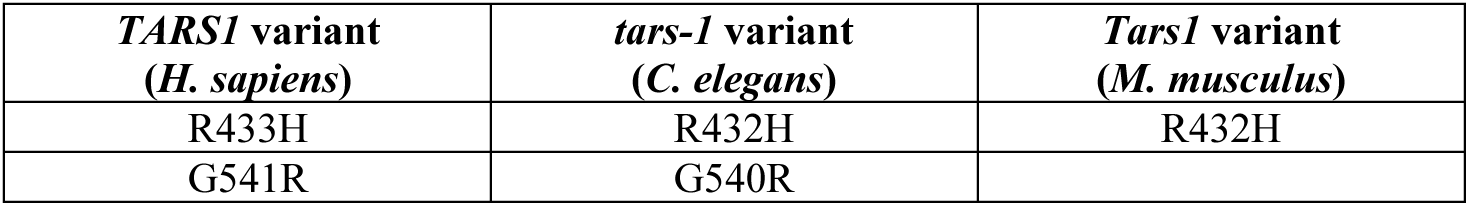
Comparison of orthologous amino-acid codons between human *TARS1*, worm *tars-1*, and mouse *Tars1*.

**Supplemental Table 2.**
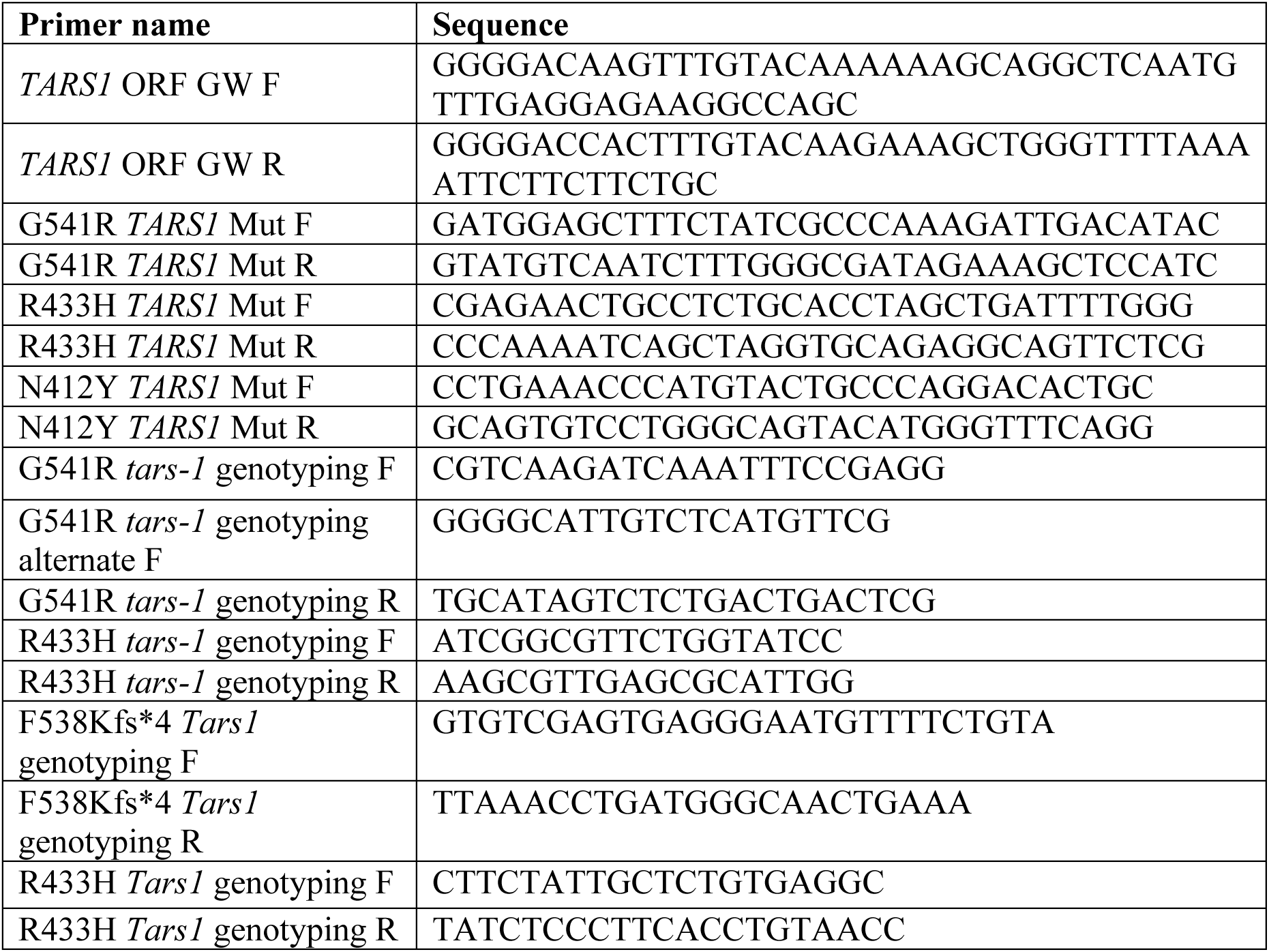
Sequences for primers used in this study. All primers are listed 5’ to 3’.

**Supplemental Table 3.**
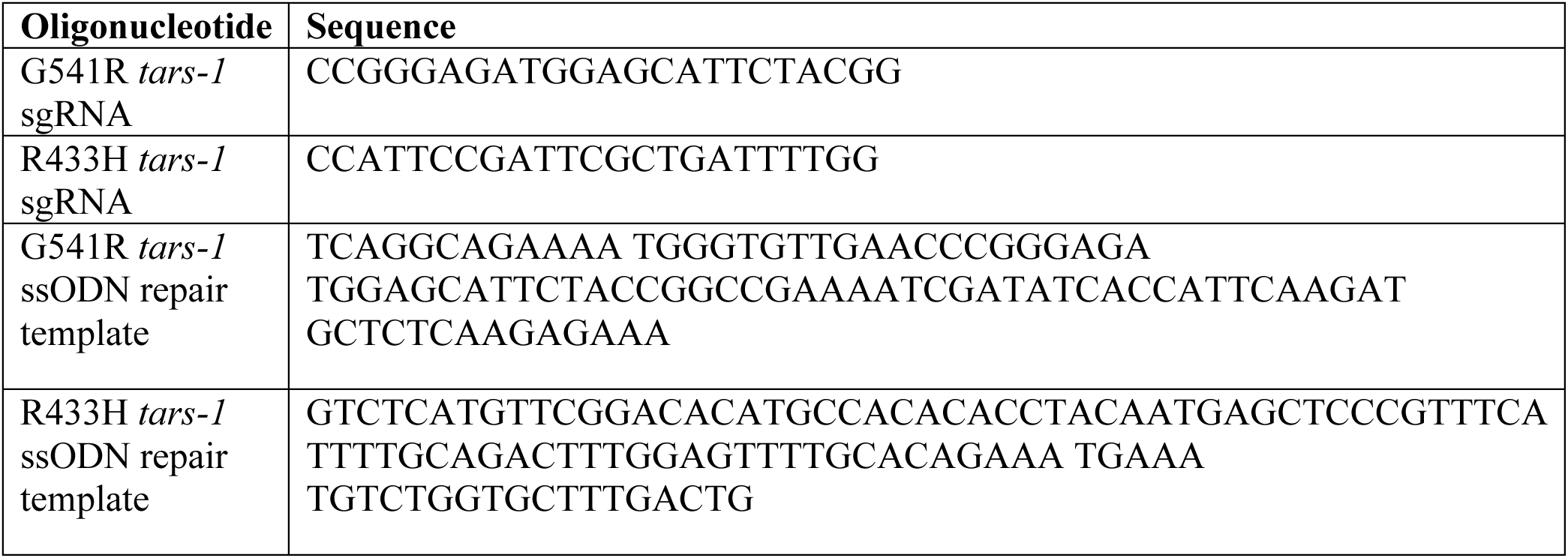
Sequences for sgRNAs and ssODNs used in this study, listed 5’ to 3’.

