## Supplemental Information for "Predictive modeling provides insight into the clinical heterogeneity associated with *TARS1* loss-of-function mutations"

**Supplemental Materials:**

| ***TARS1* variant**  **(*H. sapiens*)** | ***tars-1* variant**  **(*C. elegans*)** | ***Tars1* variant**  **(*M. musculus*)** |
| --- | --- | --- |
| R433H | R432H | R432H |
| G541R | G540R |  |

**Supplemental Table 1**. Comparison of orthologous amino-acid codons between human *TARS1*, worm *tars-1*, and mouse *Tars1.*

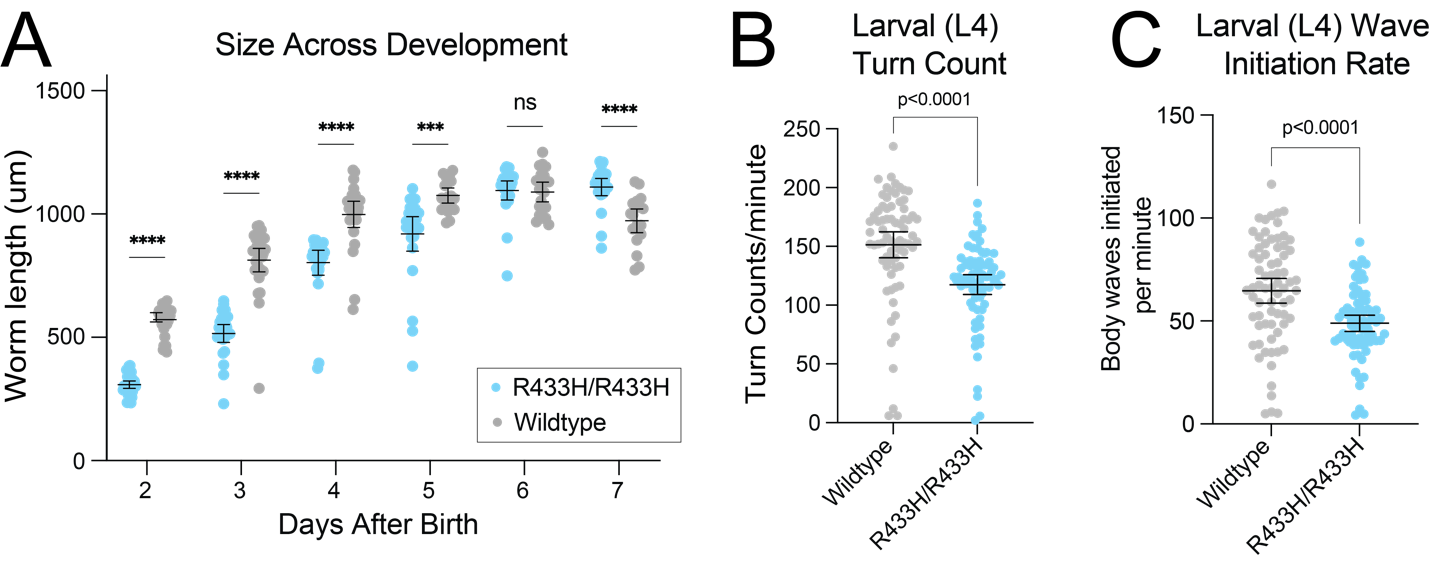

**Supplemental Figure 1**. **(A)** Body length measurements of R433H/R433H *tars-1* worms and wild-type *tars-1* worms from two to six days after birth. For R433H/R433H, 24-29 worms were measured each day. For wild-type worms, 19-35 were measured each day.  **(B)** Turn counts per minute for R433H/R433H worms (n=73) and wild-type worms (n=71) at larval stage L4, which was identified based on gonadal development. **(C)** Rate of wave initiations from either the head or the tail for L4 R433H/R433H worms (n=73) and wild-type worms (n=71). For all panels, bars indicate mean value and 95% confidence intervals. Statistical significance was evaluated using an unpaired t-test with Welch’s correction; ****, p<0.0001; ***, p<0.001 ns=not significant.

**Movie 1. R433H/R433H worms display significant locomotion defects.** A ten-second video of adult (P9) wildtype worms (N2 strain) thrashing in liquid M9 media, followed by a ten-second video clip of P9 R433H/R433H *tars-1* worms thrashing in liquid M9.

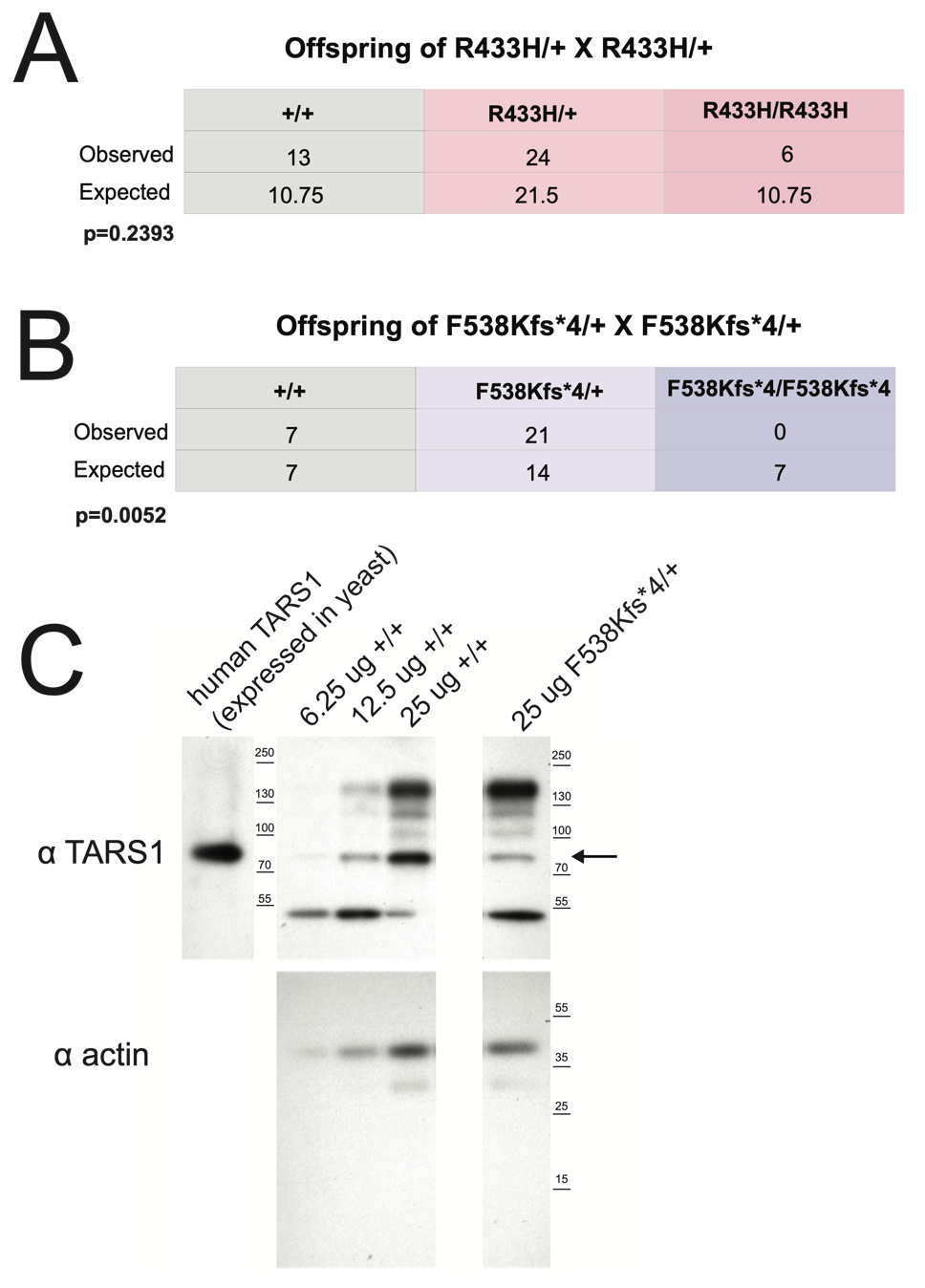

**Supplemental Figure 2**. **(A)** Genotype analysis of 28 offspring from *Tars1*^F538Kfs*4/+^ x *Tars1*^F538Kfs*4/+^ mouse mating pairs. **(B)** Genotype analysis of 43 offspring from *Tars1*^R433H/+^ x *Tars1*^R433H/+^ mouse mating pairs. All mice were genotyped at approximately 3 weeks of age. Chi-square tests were performed to determine if the difference between observed genotype counts and expected genotype counts was statistically significant. **(C)** Representative western blot image for Tars1 protein in brain tissue of wild-type *Tars1* mice and *Tars1*^F538Kfs*4/+^ mice. Human TARS1 (predicted size of 83 kDa), expressed in yeast, is shown on the left as a size control. An arrow points to the band corresponding to Tars1 (predicted size of 83 kDa). For wild-type mice, 6.25μg, 12.5μg, and 25μg lysate was loaded as a comparison to 25μg lysate for *Tars1*^F538Kfs*4/+^ samples.

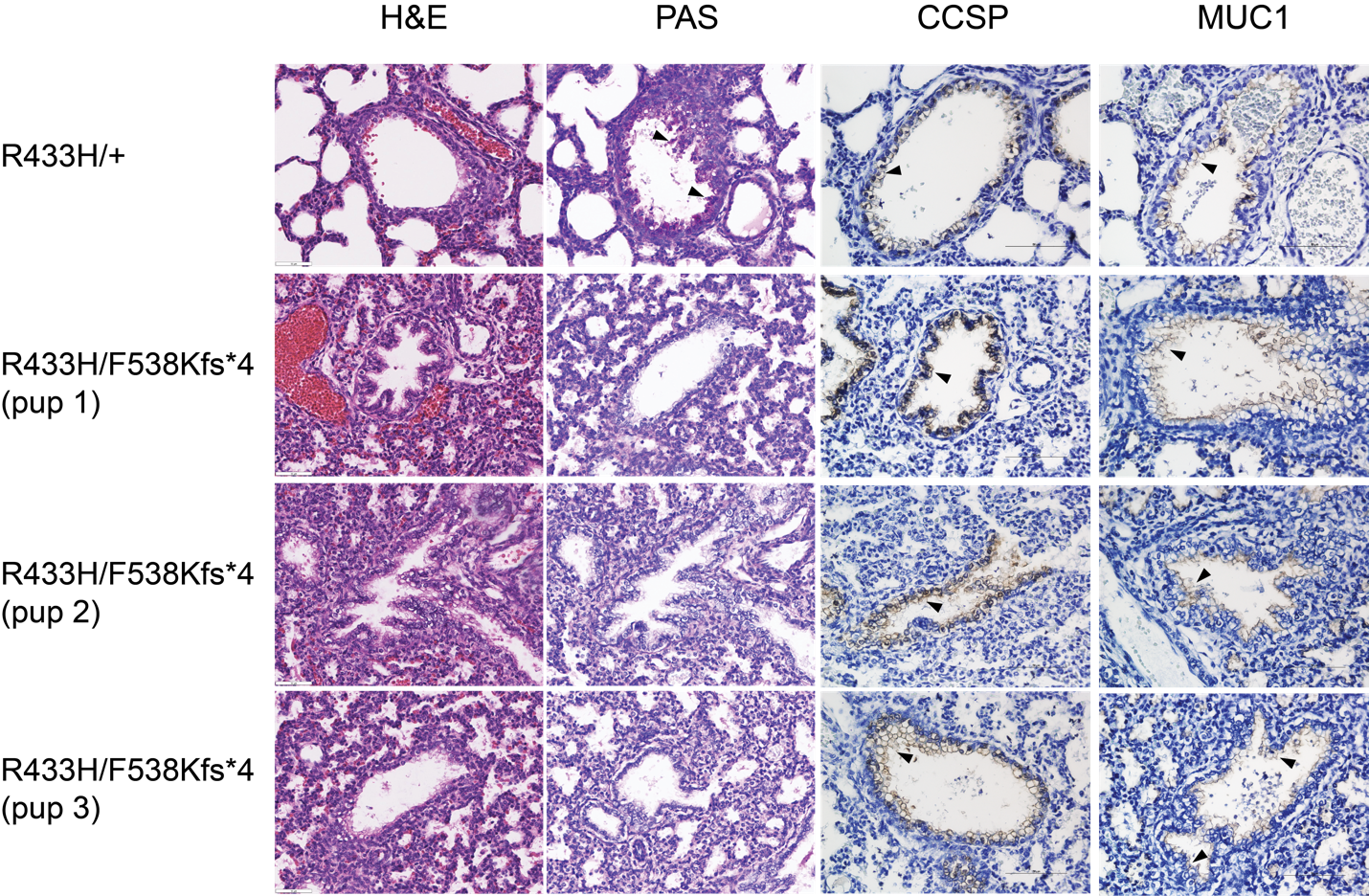
**Supplemental Figure 3.** Lung sections stained with (from left to right) H&E, PAS, CCSP antibody, or MUC1 antibody. The R433H/+ images (top row) are representative of three R433H/+ P0 pups, and the R433H/F538Kfs*4 images (bottom three rows) are representative of five R433H/F538Kfs*4 P0 pups. Black arrows point to the PAS signal in the R433H/+ mouse, and the CCSP and MUC1 signals in all mice. The CCSP and MUC1 signals are shown in brown, with the hematoxylin counterstain in dark blue.

| **Primer name** | **Sequence** |
| --- | --- |
| *TARS1* ORF GW F | GGGGACAAGTTTGTACAAAAAAGCAGGCTCAATGTTTGAGGAGAAGGCCAGC |
| *TARS1* ORF GW R | GGGGACCACTTTGTACAAGAAAGCTGGGTTTTAAAATTCTTCTTCTGC |
| G541R *TARS1* Mut F | GATGGAGCTTTCTATCGCCCAAAGATTGACATAC |
| G541R *TARS1* Mut R | GTATGTCAATCTTTGGGCGATAGAAAGCTCCATC |
| R433H *TARS1* Mut F | CGAGAACTGCCTCTGCACCTAGCTGATTTTGGG |
| R433H *TARS1* Mut R | CCCAAAATCAGCTAGGTGCAGAGGCAGTTCTCG |
| N412Y *TARS1* Mut F | CCTGAAACCCATGTACTGCCCAGGACACTGC |
| N412Y *TARS1* Mut R | GCAGTGTCCTGGGCAGTACATGGGTTTCAGG |
| G541R *tars-1* genotyping F | CGTCAAGATCAAATTTCCGAGG |
| G541R *tars-1* genotyping alternate F | GGGGCATTGTCTCATGTTCG |
| G541R *tars-1* genotyping R | TGCATAGTCTCTGACTGACTCG |
| R433H *tars-1* genotyping F | ATCGGCGTTCTGGTATCC |
| R433H *tars-1* genotyping R | AAGCGTTGAGCGCATTGG |
| F538Kfs*4 *Tars1* genotyping F | GTGTCGAGTGAGGGAATGTTTTCTGTA |
| F538Kfs*4 *Tars1* genotyping R | TTAAACCTGATGGGCAACTGAAA |
| R433H *Tars1* genotyping F | CTTCTATTGCTCTGTGAGGC |
| R433H *Tars1* genotyping R | TATCTCCCTTCACCTGTAACC |

**Supplemental Table 2**. Sequences for primers used in this study. All primers are listed 5’ to 3’.

| **Oligonucleotide** | **Sequence** |
| --- | --- |
| G541R *tars-1* sgRNA | CCGGGAGATGGAGCATTCTACGG |
| R433H *tars-1* sgRNA | CCATTCCGATTCGCTGATTTTGG |
| G541R *tars-1* ssODN repair template | TCAGGCAGAAAA TGGGTGTTGAACCCGGGAGA TGGAGCATTCTACCGGCCGAAAATCGATATCACCATTCAAGAT GCTCTCAAGAGAAA |
| R433H *tars-1* ssODN repair template | GTCTCATGTTCGGACACATGCCACACACCTACAATGAGCTCCCGTTTCA TTTTGCAGACTTTGGAGTTTTGCACAGAAA TGAAA TGTCTGGTGCTTTGACTG |

**Supplemental Table 3**. Sequences for sgRNAs and ssODNs used in this study, listed 5’ to 3’.
